# ER-located Ca^2+^ ATPase ACA2 regulates Ca^2+^ cytoplasmic pool linked to root hair growth in *Arabidopsis thaliana*

**DOI:** 10.64898/2026.07.06.736746

**Authors:** Mariana Carignani Sardoy, Vinicius Ávila Cabral, Julian Garcia Bossi, Stefano Buratti, Alessia Candeo, Giorgia Tortora, Paulina Ramirez Miranda, Cecilia Borassi, Victoria Berdion Gabarain, Javier Martinez Pacheco, Diana Rosa Rodríguez-García, Cristina Marino Buslje, Jorge P. Muschietti, Andrea Bassi, Elke Barbez, Arthur Fernandes Stradiotto Marcusse, Maria Teresa Portes, Daniel S. C. Damineli, Hugo Verli, Alex Costa, José M. Estevez

**Affiliations:** Fundación Instituto Leloir and IIBBA-CONICET. Av. Patricias Argentinas 435, Buenos Aires C1405BWE, Argentina; Universidade Federal do Rio Grande do Sul, Centro de Biotecnologia, Porto Alegre, RS, Brasil; Instituto de Investigaciones en Ingeniería Genética y Biología Molecular, Dr. Héctor Torres (INGEBI-CONICET), Vuelta de Obligado 2490, Buenos Aires, C1428ADN, Argentina; Dipartimento di Bioscienze, Università degli Studi di Milano, Milano, Italia; Dipartimento di Fisica, Politecnico di Milano, Milano, Italia; Institute of Biology II, Molecular Plant Physiology (MoPP) University of Freiburg, Germany. Center for Integrative Biological Signalling Studies (CIBSS) University of Freiburg, Germany and Spemann Graduate School of Biology and Medicine (SGBM), University of Freiburg, Germany; Laboratory of Biochemistry, Wageningen University, The Netherlands; Departamento de Biodiversidad y Biología Experimental, Facultad de Ciencias Exactas y Naturales, Universidad de Buenos Aires, Int. Güiraldes 2160, Ciudad Universitaria, Pabellón II, C1428EGA Buenos Aires, Argentina; Department of Botany, Institute of Biosciences, University of São Paulo, São Paulo, Brazil; Center of Mathematics, Computation and Cognition, Federal University of ABC, São Bernardo, São Paulo, Brazil; Brazilian Biosciences National Laboratory (LNBio), Brazilian Center for Research in Energy and Materials (CNPEM), Campinas 13083-100, SP, Brazil; Istituto di Biofisica, Consiglio Nazionale delle Ricerche (CNR), Milano, Italia; ANID - Millennium Science Initiative Program - Millennium Institute for Integrative Biology (iBio) Santiago, Chile; Centro de Biotecnología Vegetal, Facultad de Ciencias de la Vida, Universidad Andrés Bello, Santiago, Chile

**Keywords:** Arabidopsis, root hair, ACA pumps, calcium, calcium oscillations, light sheet fluorescence microscopy, YC3.6

## Abstract

Root hairs (RH) are excellent model systems for studying cell size and polarity since they elongate several hundred-fold their original size. Their tip growth is regulated by both intrinsic and environmental signals and is associated with the existence of a highly controlled cytoplasmic tip Ca²⁺ gradient, whose disruption impairs RH development. The molecular mechanisms underlying the Ca^2+^ homeostasis fine tuning and the Ca^2+^ organellar contributions to the cytoplasmic pool remain unclear. In the model plant *Arabidopsis thaliana*, many efflux routes are present, including those that employ Ca^2+^-pumps from the Autoinhibited Ca^2+^-ATPase (ACA) family. Here, we identified that the ER localized ACA2, and to a lower extent ACA7, are crucial ACAs required to control RH growth. By using genetically encoded Ca^2+^ biosensors we showed that Ca^2+^-dynamics are compromised in the *aca2-2* mutant, having lower cytosolic Ca^2+^ concentration [Ca^2+^]_cyt_ and growth rate, showing an altered homeostatic calcium setpoint compared to Col-0. Accordingly, the ACA2 mutation changed the dynamics of [Ca^2+^]_cyt_ oscillations coupled to growth rate, inducing longer periods and more regular oscillations in the dominant high-frequency range (around 22 s), and slower oscillations (around 1 min) in the low-frequency range. Finally, expression of ACA2 with changes in four putative Ca^2+^ binding residues (ACA2ΔCa^2+^) failed to rescue the RH growth phenotype in the *aca2-2* mutant. Collectively, our findings indicate that ER-localized ACA2 and possibly ACA7 are crucial for modulating cytoplasmic Ca^2+^ signals, possibly composing a critical part of a negative feedback loop, and their absence leads to impairments in RH cell elongation.

## Introduction

Plant roots are a subject of interest in biology for their role as plant anchors and because they oversee water and nutrient intake. Root hairs (RH), also known as trichoblasts, are fundamental for plant development as they increase root surface area for water and nutrient absorption and improve anchoring to the ground, as well as being the infection site for nitrogen-fixing bacteria. RH presents a specialized type of growth known as polar or tip growth. In this process there is a highly controlled cell wall remodeling that allows RH to withstand the turgor pressure forces acting during growth (Balcerowicz et al., 2015; Lopez et al., 2024; Ryan et al., 2001). RH growth is oscillatory with alternating higher and lower growth rates. Polar cell growth is sustained by oscillatory feedback loops comprising three main components that, together, play an important role in regulating this process although their fine tune regulation is not fully understood. One of the main components are Reactive Oxygen Species (ROS) that, together with calcium ions (Ca^2+^) and pH, maintain polar growth over time. Ca^2+^ is involved in practically all facets of plant development, including responses to both biotic and abiotic factors (Kudla et al., 2018). Fluctuations in cytosolic Ca^2+^ concentration [Ca^2+^]_cyt_ arise from the synchronized regulation of influx and efflux pathways, with distinct Ca^2+^ circuits operating in various subcellular locations (Aldon et al., 2018; Corso et al., 2018; García Bossi et al., 2020; Johnson et al., 2019; Kudla et al., 2018; Martí Ruiz et al., 2020; Moeder et al., 2019; Resentini et al., 2021; Tang & Luan, 2017). High levels of free [Ca^2+^]_cyt_ are cytotoxic, which is why it must be kept at submicromolar concentrations (Clapham, 2007). For this reason, the increase in [Ca^2+^]_cyt_ concentration during signaling events is followed by storage of Ca^2+^ into cellular compartments or export to the apoplast although the underlying mechanisms remain unclear (Bonza et al., 2013; Costa et al., 2018, 2023; Sanders et al., 1999; White & Broadley, 2003; Grenzi et al., 2025).

Ca^2+^ moves from the extracellular space and intracellular reservoirs to the cytoplasm through Ca^2+^ permeable channels. It is then removed from the cytoplasm by Ca^2+^ transporters, like exchangers and pumps, and sent back to extracellular space and organelles (Demidchik et al., 2018). Within this group of transporters, there are P-type IIA Ca^2+^ ATPases (ECAs), analogous to animal sarcoplasmic-endoplasmic reticulum Ca^2+^ ATPases (SERCA), and P-type IIB Ca^2+^ ATPases (ACAs), which are equivalent to animal PM-type ATPases (PMCA) (Axelsen & Palmgren, 2001; Baxter et al., 2003; Costa et al., 2023; García Bossi et al., 2020; Hocking et al., 2017; Ma & Berkowitz, 2017; Pittman & Hirschi, 2016). ACA-type pumps are regulated by the binding of Ca^2+^/calmodulin to an N-terminal regulatory domain, which alleviates autoinhibition and subsequently activates the pump. This establishes a feedback loop where the Ca^2+^ influx produces a “transient” signal that subsequently activates ACA-efflux pathways to restore Ca^2+^ to resting basal levels. Located in the cytoplasm, ACAs present 3 characteristic domains: actuator (A domain), nucleotide binding (N domain) and phosphorylation (P domain). Meanwhile, the transmembrane domains are described as transport domain (T domain), where the Ca^2+^ binding site is, and support domain (S domain) (Palmgren & Nissen, 2011).

Ca^2+^ pumps and exchangers have been identified in several membrane systems, including the endoplasmic reticulum, vacuole, and plasma membrane. Although ACAs, ECAs, and cation/proton exchangers (CAXs) remove Ca^2+^ from the cytosol, their distinct roles in modulating the intensity or duration of various stimulus-specific Ca^2+^ signals remain only superficially comprehended (Baekgaard et al., 2005; Bonza et al., 2000; Chung et al., 2000; Costa et al., 2017, 2023; Giacometti et al., 2012; Wang et al., 2024). There are 10 different ACAs in *Arabidopsis thaliana* (García Bossi et al., 2020) and few of them have been fully characterized. The plasma membrane-localized ACAs (ACA8,9,10,12,13) have been associated with plant development (George et al., 2008; J. Zhang et al., 2014), compromised reproduction (Iwano et al., 2014; Schiøtt et al., 2004; Yu et al., 2018), and inadequate responses to pathogens (for ACA8 and ACA10; (Frey et al., 2012)). ACA4 and ACA11 are directed to vacuoles (Hilleary et al., 2020; Lee et al., 2007) and exhibit a role in salicylic acid (SA)-dependent lesions in leaves (Boursiac et al., 2010). In a recent study, the vacuolar ACA4 and ACA11 and, to a lower extent, the ER-localized ACA2 and ACA7, were all reported to be important in the regulation of Ca^2+^ dynamics linked to pollen tube growth (García Bossi et al., 2026). Our work presents evidence that the ER-localized ACA2 and ACA7 are functionally relevant for RH growth. Growing *aca2* RH has a lower [Ca^2+^]_cyt_ homeostatic setpoint and abnormal oscillations coupled to growth. These findings highlight the importance of the ER-localized ACA2 in regulating cellular Ca^2+^ dynamics, while the absence of their activities impair RH growth.

## Results and Discussion

To determine which ACAs are expressed in RH cells and might affect the Ca^2+^_cyt_ dynamics during cell elongation, we performed an *in-silico* analysis using Arabidopsis RootCellAtlas tool (**Figure 1A** and **Figure S1**). This atlas integrates the information from single cell RNAseq data (Denyer et al., 2019; Jean-Baptiste et al., 2019; Ryu et al., 2019; Shahan et al., 2022; Shulse et al., 2019; Wendrich et al., 2020; T.-Q. Zhang et al., 2019) and can identify cell types in 2D uniform manifold approximation and projection (UMAP). The expression of the 10 Arabidopsis ACA genes in RH cells, including atrichoblasts and trichoblasts, in agreement with previous analysis (García Bossi et al., 2020) showed that *ACA2*, *ACA7* and *ACA11* are highly expressed in the root epidermis (**Figure 1B-C** and **Figure S1**) showing a preferential expression in trichoblasts. As for other ACAs, these genes present low levels of expression in the epidermis (**Figure S1**). To examine the roles of the ACA genes in RH growth, two previously published T-DNA insertion mutants (Lucca & León, 2012; Rahmati Ishka et al., 2021) were analysed for each *ACA* and a double mutant was also generated (**Figure 1D**). We analysed RH growth in plants growing *in vitro* (**Figure 1E-F**). RH growth is drastically affected in *aca2-1* and *aca2-2*, and to a lower extent, in *aca7-1* and *aca7-2* mutants while the double mutant *aca2-2/aca7-1* shows no significant differences from *aca2-1* mutant (**Figure 1E-F)**. It is surprising that the double mutant aca2-2/aca7-1 exhibited a minor rescue in the RH phenotype compared to aca2-2, indicating a potential realignment at either the transcriptional or post-translational level in the remaining non-mutated ACAs. Alternatively, given that the disruption of ER- and vacuolar-localized ACAs has been linked to the activation of salicylic acid (SA)-dependent responses (Boursiac et al., 2010; Rahmati Ishka et al., 2021), we cannot rule out that the partial recovery observed in the *aca2-2/aca7-1* double mutant involves a low-level SA response; this potential connection between ER-ACA function, SA signaling and RH growth deserves further investigation. Then, we studied plants growing in soil (**Figure 1G-H**) using rhizoboxes to analyse root growth in conditions that more accurately resemble the environment in which plants grow in nature. In contrast to *in vitro* growth in plates, by developing in soil, the roots were exposed to less light and were able to interact with microorganisms. In these conditions the restricted RH growth was still significant in *aca2-2*, whereas *aca7-1* was not significantly different from Col-0 (**Figure 1G**). Both *aca4* and *aca11* mutants did not show significant differences in RH phenotype when compared to Col-0 (**Figure S1**). However, the *aca4/aca11* double mutant did show a reduced RH growth (**Figure S1D**). This result indicates that ACA2, and ACA7 to a lower extent, are both required for proper RH growth.

**Figure 1.**
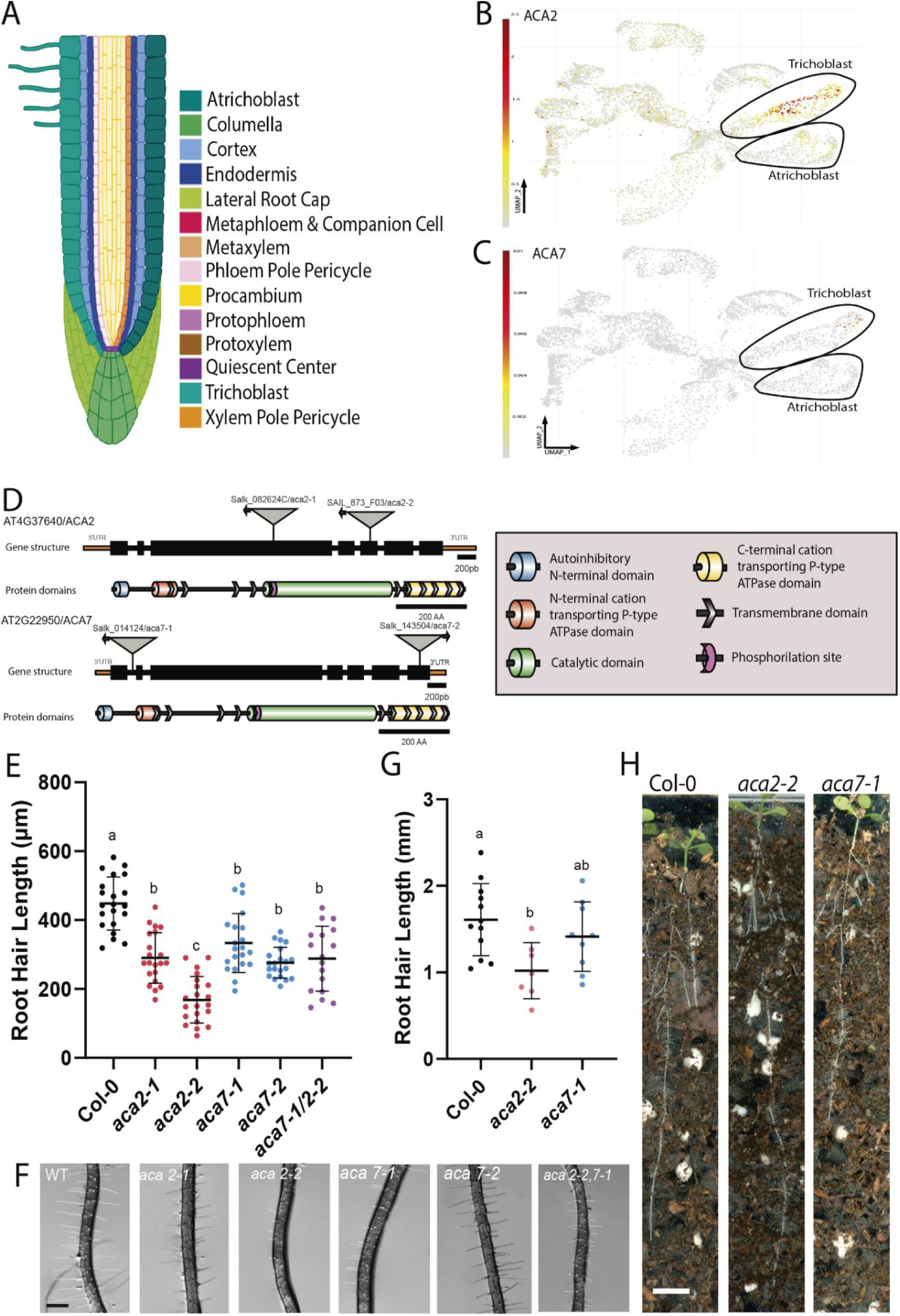
Role of ACA2 and ACA7 in RH growth. (**A**) Schematic illustration of root (right) and (B-C) UMAP showing expression of ER-ACA2 (**B**) and ER-ACA7 (**C**) in trichoblast and atrichoblast cells. UMAPs showing expression levels of *ACA2* and *ACA7* are normalized to marker genes’ expression. These UMAPS were generated by RootCellAtlas initiative 2022 (https://rootcellatlas.org/). (**D**) Diagrams for *ACA2* and *ACA7* genes including T-DNA insertions and their respective protein domains. Gene structure and T-DNA insertion position were based on Araport11 genome information available on TAIR (Berardini et al., 2015). Protein domain was compiled from Uniprot (UniProt Consortium, 2023) and Pfam (now InterPro) (Paysan-Lafosse et al., 2023) databases. Diagrams were made using the IBS 2.0 Web Server (Xie et al., 2022). (**E**) Scatterplot of RH length of Col-0, *aca2-1, aca2-2, aca7-1, aca7-2* and *aca2-2/aca7-1* mutants grown *in vitro* at 22°C. Each point is the mean of the length of the 10 longest RHs identified in the maturation zone of a single root. Data is the mean ± SD, n = 17-21 roots, statistical analysis was performed using a Generalized Least Squares (GLS) model. Post hoc comparisons were conducted using Tukey’s test to assess differences between groups. Different letters indicate statistically significant differences (p < 0.05). (**F**) Representative images of each genotype quantified in E. Scale bar = 200 µm. (G-H) Phenotype analysis in Rhizobox, a growth condition in soil that is distinct from, and not redundant with, the in vitro assay shown in (E). (**G**) Scatterplot of RH length for WT Col-0, *aca2-2*, and *aca7-1* plants. Each point represents the average hair length of a root in the elongation zone; 10 root hairs were measured per plant. Error bar represents mean ± standard deviation, n = 7-12 roots. Statistical analysis: A Generalized Least Squares (GLS) model was used. For post hoc comparisons, a Tukey test was conducted; groups marked with different characters have significant differences between them (p < 0.05). (**H**) Representative images of each genotype presented in the graph in G. Scale bar = 500 µm.

Considering these findings, we investigated the impact of *aca2-2* mutant on Ca^2+^ dynamics and, subsequently, in RH growth. For this, we performed light sheet fluorescence microscopy (LSFM) imaging with plants expressing the Förster Resonance Energy Transfer (FRET) Yellow Cameleon (NES-YC3.6) sensor (Krebs et al., 2012; Nagai et al., 2004) (**Figure 2A-B**). This technique allows a rapid acquisition of images with a high dynamic range with single-cell resolution over a wide field of view for vertically growing seedlings (Berthet & Maizel, 2016; Costa et al., 2013). In this condition, RH can develop for extended time under near-physiological circumstances (Candeo et al., 2017) and this allowed us to record FRET in actively growing RH and estimate the Ca^2+^ oscillation frequency at the tip. We monitored RH development under low light exposure at the root maturation zone, where the growth and elongation of both newly formed and mature RH were monitored (**Supplementary Movie S1**). Growing RHs were manually chosen, and their growth over time tracked with image registration, allowing the extraction of the FRET signal from a designated region of interest (ROI) at the apex of the RH. Tip [Ca^2+^]_cyt_ oscillations were assessed by the FRET ratio through time for each RH (**Figure 2C-D, Figure S2**). The ratiometric biosensor indicated a lower concentration of cytosolic Ca^2+^ in *aca2-2* (**Figure 2E**). This may seem counterintuitive, since ACA2 would be pumping Ca^2+^ into the ER, removing it from the cytosol. However, the regulation of cytosolic Ca^2+^ concentration relies heavily on ER stores, thus a decrease in Ca^2+^ reservoir and altered time scale can indeed lead to a lower [Ca^2+^]_cyt_ (Xiong et al., 2025). Thus, the mutation in ACA2 changes Ca^2+^ regulation across intracellular compartments, effectively lowering the Ca^2+^ homeostatic setpoint. In addition, a reduced mean growth rate was found in the *aca2-2* in this LSFM growth condition (**Figure 2F**), being consistent with the final RH length (**Figure 1**). This can decrease the tip-focus gradient and lead to a premature growth arrest, akin to the effect seen in H^+^-pumps mutants in pollen tubes (Hoffmann et al., 2020). Changing such an integrated control system may impact the entire dynamics between Ca^2+^ and growth, warranting an investigation of their oscillations.

**Figure 2.**
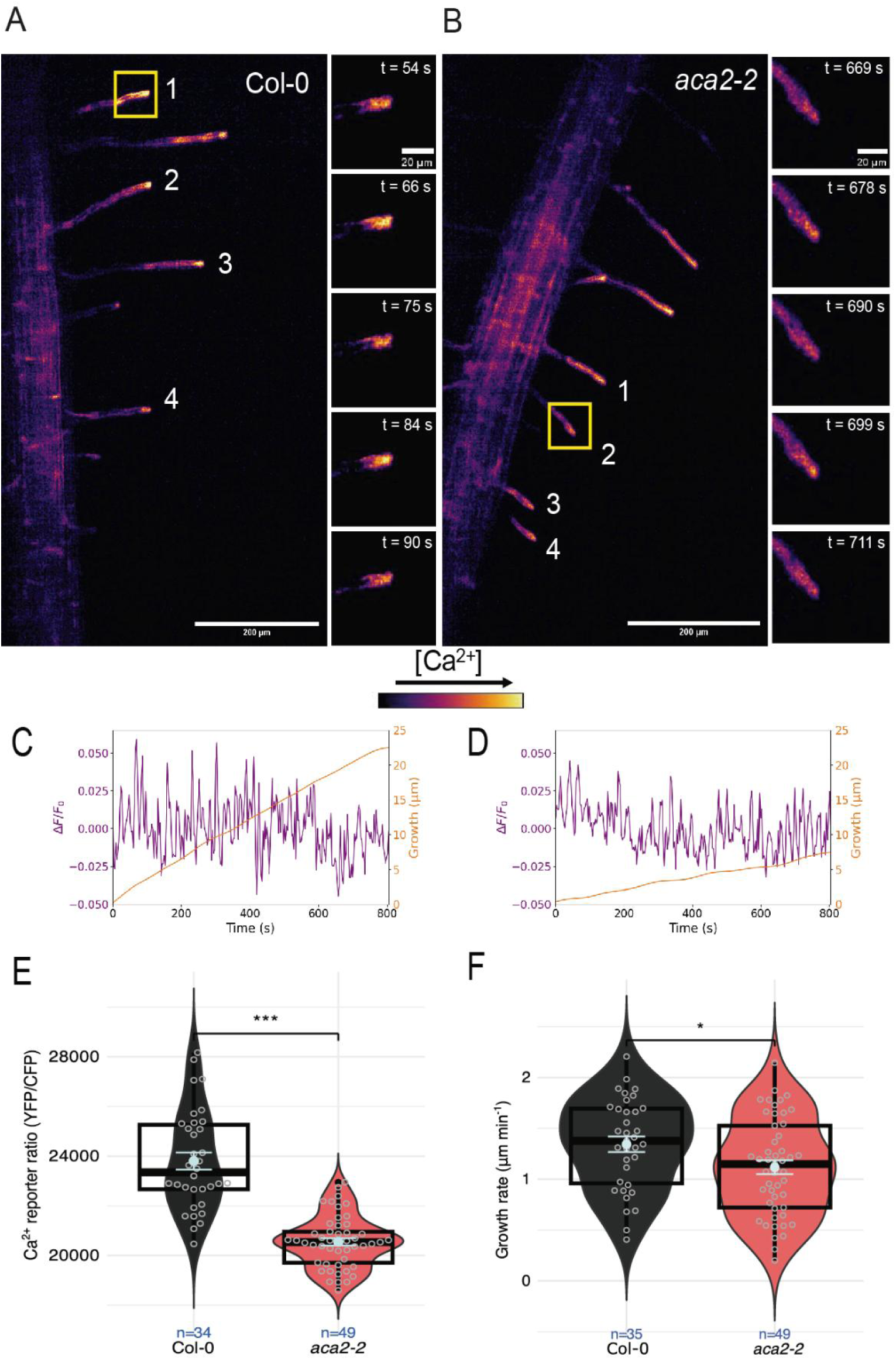
Analysis of Ca^2+^ dynamics using NES-YC3.6 biosensor. (**A-B**) Maximum Intensity Projection (MIP) of the ratio between the cpVenus and CFP fluorescent channels images obtained from 12 slices of the sample spaced 4 μm apart for Col-0 (A) and *aca2-2* (B). On the right we show the MIP of a root hair of the ratio between cpVenus and CFP fluorescent channels at different time points for Col-0 and *aca2-2*. (**C-D**) Representative oscillations of FRET signal in a root hair (purple) and growth of the measured RH (orange) in Col-0 (**C**) and *aca2-2* (**D**). (**E**) [Ca^2+^]_cyt_ estimated with a ratiometric biosensor (YC3.6) measured at the RH tip. Violin plots show kernel density, individual observations as grey dots, with boxplots and overlayed mean ± SEM (light blue) summarizing the central tendency and dispersion. Statistical comparison between groups was determined by a t-test (***, p < 0.001). (**F**) RH growth rate observed during measurement of oscillations. Violin plots show kernel density, individual observations as grey dots, with boxplots and overlayed mean ± SEM (light blue) summarizing the central tendency and dispersion. Statistical comparison between groups was determined by a t-test (*, p < 0.05).

An overall Fourier analysis of RH [Ca^2+^]_cyt_ oscillations revealed potential differences in both high-(HF; from 0.05 to 0.028 Hz or periods from 20 to 36 s) and low-frequency (LF; from 0.013 to 0.003 Hz or 1.3 min to 5 min) regimes of [Ca^2+^]_cyt_ oscillations (**Figure S3**). These two distinct oscillatory components have been reported (Brost et al., 2019; Monshausen et al., 2008), with HF oscillations associated with the pulsatile RH growth (Candeo et al., 2017). To analyze the HF [Ca^2+^]_cyt_ growth rate, sufficient resolution of the imaging and growth rate, sufficient resolution of the imaging and tracking technique are required, while for LF a long acquisition time span is critical. However, our dataset obtained with a 10x objective lens was significantly influenced by noise in the HF range. Thus, we increased the magnification to 20x and extended the acquisition time specifically to assess [Ca^2+^]_cyt_ tracking technique is required growth rate, sufficient resolution of the imaging and tracking technique are required, while for LF a long acquisition time span is critical. However, our dataset obtained with a 10x objective lens was significantly influenced by noise in the HF range. Thus, we increased the magnification to 20x and extended the acquisition time specifically to assess [Ca^2+^]_cyt_, while for LF a long acquisition time span is critical. However, our dataset obtained with a 10x objective lens was significantly influenced by noise in the HF range. Thus, we increased the magnification to 20x and extended the acquisition time specifically to assess [Cagrowth rate, sufficient resolution of the imaging and tracking technique are required, while for LF a long acquisition time span is critical. However, our dataset obtained with a 10x objective lens was significantly influenced by noise in the HF range. Thus, we increased the magnification to 20x and extended the acquisition time specifically to assess [Ca^2+^]_cyt_ oscillations synchronized with growth rate. The time series of both [Ca^2+^]_cyt_ and growth rate with increased resolution were detrended (**Figure 3A-B**), and their synchronized oscillations analyzed with a cross-wavelet transform (**Figure 3C-D**). The significant joint periodicity occurring between [Ca^2+^]_cyt_ and growth rate was obtained for each time point, showing the prevalence of oscillations in the high frequency range, but with the clear presence of low frequency components (see the power spectrum around 2 min in **Figure 3C-D**). Thus, we divided the analysis focusing on HF and LF components separately, which allowed detecting a higher mean period in HF oscillations (**Figure 3E**). The main periodic components were estimated throughout time by the wavelet ridges, which are continuous changes in the power peaks depicted with grey traces (**Figure 3C-D**) within the significant regions (surrounded by white) of the wavelet spectrum. There is clear variation in the periodic components through time (non-stationarity), with some series being more regular than others. We found that *aca2-2* RHs have more regular synchronized oscillations between [Ca^2+^]_cyt_ and growth rate in the HF component, indicated by a lower coefficient of variation in wavelet ridges (**Figure 3F**) and lower dispersion of phase relationships (**Figure S4**). Such increased regularity can be seen in the representative time series as a more continuous and flatter wavelet ridge in *aca2-2*, with more coherent phase arrows (**Figure 3C-D**). As for LF oscillations, the mutant showed a significantly lower period of synchronized [Ca^2+^]_cyt_/growth oscillations (**Figure 3G**). Both phenotypes were already indicated by the overall comparisons of Fourier PSDs of [Ca^2+^]_cyt_ oscillations in the 10x dataset (**Figure S3**). We further validated these phenotypes with wavelet analyses of the individual [Ca^2+^]_cyt_ in both datasets (**Figure S5**). This comprehensive assessment showed coherent results, with the same tendency even in the few cases where statistical significance was not found, supporting the interpretation of a robust phenotype.

**Figure 3.**
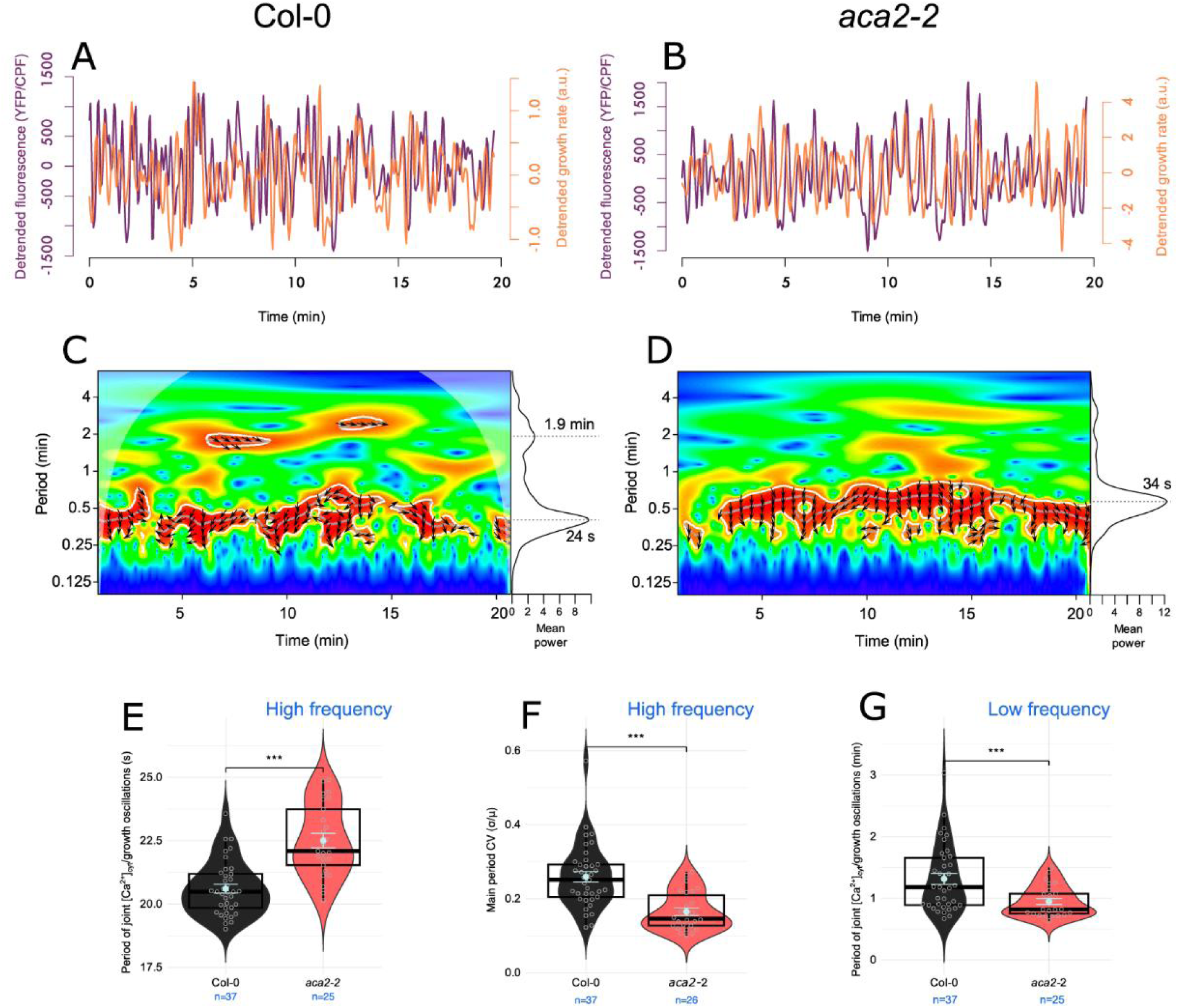
Synchronized oscillations between [Ca^2+^]_cyt_ and growth rate are altered in the *aca2-2* mutant. (**A–B**) Representative detrended time series of cytosolic Ca^2+^ biosensor and growth rate in Col-0 (**A**) and *aca2-2* (**B**). Purple traces show the fluorescence ratio of the biosensor YC3.6, reflecting oscillatory changes in [Ca^2+^]_cyt_. Orange traces show the corresponding growth rate series. Detrending removes baseline changes to focus on biologically meaningful oscillatory dynamics. (**C–D**) Significant joint periodicity of [Ca^2+^]_cyt_ and growth rate evaluated with a cross-wavelet transform for Col-0 (**C**) and *aca2-2* (**D**). Heat colors represent joint wavelet power, where warmer colors indicate higher power. Panels to the right of the wavelet spectrum show the mean wavelet power across time, with labeled peaks. Significance of joint periodic components was assessed with AR(1), an autoregressive process of order 1, as null model with significant regions outlined in white (p < 0.05). Main periods are shown as gray ridges, which trace continuous peaks of significant joint power through time. The phase relationship between [Ca^2+^]_cyt_ and growth rate are shown as arrows pointing towards an imaginary unit circle, with oscillations being completely in-phase to the right (0 phase difference) and in anti-phase to the left (phase difference of π), with phase differences being: 0 to π/2 (1st quadrant), π/2 to π (2nd quadrant), -π to -π/2 (3rd quadrant) and -π/2 to 0 (4th quadrant). Thus, the downward direction denotes growth rate leading Ca^2+^ oscillations, which predominates, and the reverse for the upward direction. (**E**) Significant periods of synchronized oscillations between [Ca^2+^]_cyt_ and growth rate, extracted from cross-wavelet ridges in the high-frequency range for each time series (grey dots are the individual mean). Violin plots show kernel density, boxplots with overlayed mean ± SEM (light blue) summarize the central tendency and dispersion, with statistical comparison between groups determined by a t-test (***, p < 0.001). (**F**) Variability of the dominant periodic component of synchronized [Ca^2+^]_cyt_/growth oscillations in the high-frequency range, quantified as the coefficient of variation (CV = SD/mean) of each individual ridge. (**G**) Low frequency significant periodicity of synchronized oscillations between [Ca^2+^]_cyt_ and growth rate, calculated as in (**E**).

The longer periods of HF [Ca^2+^]_cyt_ oscillations in *aca2-2* are consistent with the substrate-depletion models (Xiong et al., 2025). In the Goldbeter–Dupont–Berridge model (Goldbeter et al., 1990), the interval between Ca^2+^ peaks in the cytosol is limited by the rate of replenishment of Ca^2+^ stores, in this case ACA2 restoring ER calcium pools. In the case of LF [Ca^2+^]_cyt_ oscillations, a different mechanism is likely involved. Thus, increasing the timescale and decreasing the amount of Ca^2+^ flux to ER stores could explain longer periods of HF Ca^2+^ oscillations, as well as the lower cytosolic Ca^2+^ concentration, since ER concentration in *aca2-2* might be further from the activation threshold to pump out Ca^2+^ (Xiong et al., 2025). This would implicate ACA2 in the feedback loop that regulates Ca^2+^ increases in the cytosol, increasing the speed of sequestration. Changing such an integrated control system might have led to a new and more stable dynamical regime in the mutant, such as a limit cycle. This “simplified dynamics” may be more rigid and less sensitive to external cues, potentially leading to premature growth arrest by failing to respond to spatial challenges efficiently, together with a shallower tip-focused [Ca^2+^]_cyt_ gradient. To further test if this “changed” dynamics of [Ca^2+^]_cyt_may impact on the Ca^2+^ in the extracellular compartments and its interactions with the cell walls, RH were stained with ALEXA COS-488 which depicts pectin charge in the RH cell walls. The double mutant *aca2-2/aca7-1* of fluorescence, suggesting a higher degree of Ca^2+^ interacting pectins in their cell walls (**Figure S6**) also linked with delayed cell expansion dynamics (Borassi et al., 2021) while no change were detected in the single mutants *aca2-2* and *aca7-1*. We note that, because the change in growth rate can precede the change in [Ca^2+^]_cyt_, an alternative and non-mutually exclusive interpretation is that cell-wall dynamics drive the cytosolic Ca^2+^ changes rather than the reverse. In line with this, the altered distribution of pectin methyl-esterification could also reflect additional wall-associated cues, such as RALF peptides, which bind de-esterified pectin and modulate cell-wall remodeling during growth (Liu et al., 2024).

Then, we tested if *aca2-2* can be rescued by the overexpression of ACA2-GFP and this was confirmed for four independent lines, where at least 80–90% of the total RH length was recovered (**Figure 4A-B**). ER localization has been established for ACA2 (Dunkley, 2006; Hong et al., 1999; Rahmati Ishka et al., 2021). With this in mind, we identified a putative ER retention signal at _906_KGKAM_911_ in ACA2 sequence (**Figure S7**). Then, we tested if this putative ER retention signal is functional in ACA2 in a transient experiment in *Nicotiana benthamiana*. By colocalization analysis with an ER-marker (ER-KDEL-mCherry), ACA2-GFP showed a high degree (close to 0.8 Manders Coefficient) of overlap in the ER compartment (**Figure 4D**). When a modified variant of ACA2, where the two Lys of the ER retention signal were changed to Ala as _906_AGAAM_911,_ was generated (ACA2ΔER), most of the ER localization was still present in the ACA2ΔER with a similar degree of ER-colocalization. This indicates that these two Lys residues are not controlling the ER targeting of ACA2 (**Figure 4E, Figure S8**). Finally, we wanted to test the impact of selected Ca^2+^ binding residues in ACA2 in the restoration of RH growth in the *aca2-2* mutant. For that, using homology modeling coupled to AlphaFold prediction, four residues Val_407_, Glu_412_, Asn_849_, Asp_853_ were identified to be important in the Ca^2+^ binding according to the structural model generated (**Figure 4C**). When changing all four residues to Ala (ACA2ΔCa^2+^), this variant of ACA2 could not complement the *aca2-2* mutant (**Figure 4F-H**) although it still localizes to the ER (**Figure 4D-E Figure S8**). This may imply that putative Ca^2+^ binding in ACA2 protein might be crucial for ACA2 activity linked to RH growth.

**Figure 4.**
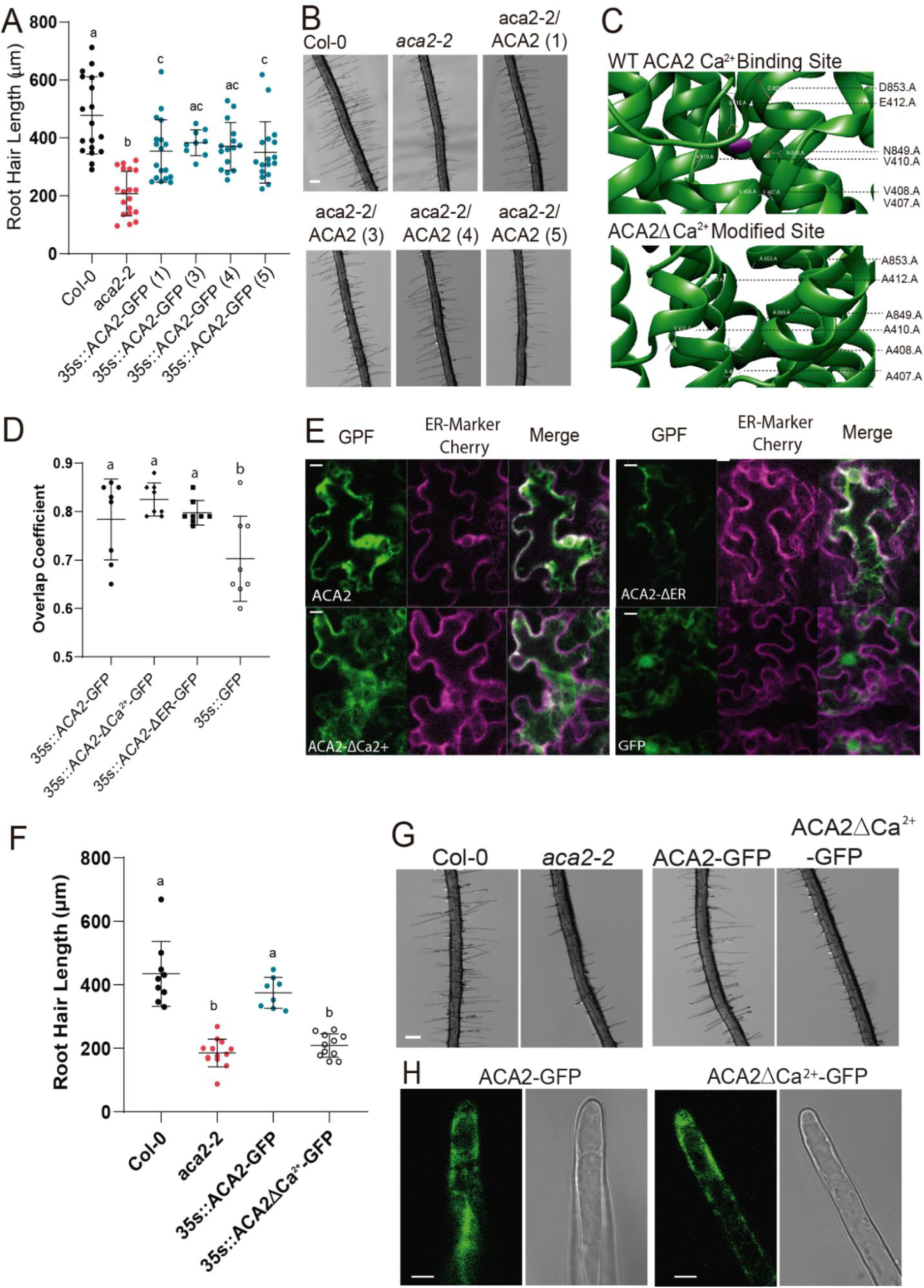
ACA2 structural features linked to RH growth functions. (**A**) Scatterplot of RH length of Col-0, *aca2-2,* and ACA2-GFP overexpressing lines (in the *aca2-2* mutant background, i.e. complementation lines) grown at 22°C. Each point is the mean of the length of the 10 longest RHs identified in the maturation zone of a single root. Data is the mean ± SD, n = 10-19 roots, statistical analysis was performed using a Generalized Least Squares (GLS) model. Post hoc comparisons were conducted using Tukey’s test to assess differences between groups. Different letters indicate statistically significant differences (p < 0.05). (**B**) Representative images of each genotype quantified in A. Scale bar = 200 µm. (**C**) Representative diagram of ACA2 and ACA2-ΔCa^2+^ Ca^2+^ binding site. (**D**) Scatterplot of Manders Coefficient between GFP and mCherry channels in confocal images. Data are the mean ± SD, n = 8, statistical analysis: One way ANOVA followed by Dunnett’s multiple comparison test. Different letters indicate statistically significant differences (p < 0.05). (**E**) Representative images of each genotype quantified in D. Scale bar = 10 µm. (**F**) Scatterplot of RH length of Col-0, *aca2-2,* and ACA2-GFP/*aca2-2* and ACA2-ΔCa^2+^-GFP/*aca2-2* overexpressing lines grown at 22°C. Each point is the mean of the length of the 10 longest RHs identified in the maturation zone of a single root. Data are the mean ± SD, n = 10-13 roots, statistical analysis was performed using a Generalized Least Squares (GLS) model. Post hoc comparisons were conducted using Tukey’s test to assess differences between groups. Different letters indicate statistically significant differences (p < 0.05). (**G**) Representative images of each genotype quantified in F. Scale bar = 200 µm. (**H)** Representative images of GFP fluorescence in RH in overexpressing lines shown in F. Scale bar = 10 µm.

To explore in more detail the structural determinant of ACA2 Ca^2+^ transport, three ACA2 variants were generated using AlphaFold3 (Abramson et al., 2024) (**Figure 5A-D**): (1) ACA2 WT: the unmodified protein, with one Ca^2+^ ion placed inside the pore and coordinated by residues V407, E412, N849, and D853 (**Figure 5A,C**); (2) ACA2 WT 2Ca^2+^: the same structure as ACA2 with an additional Ca^2+^ ion placed in the phosphorylation motif DKTGT (**Figure 5A**); (3) ACA2ΔCa^2+^ (VEND-AAAA): ACA2 in which the Ca^2+^–binding site VEND was substituted with AAAA (**Fig. 5A,C**). By molecular dynamics we tested the impact of these modifications in the ACA2 capacity of Ca^2+^ transport (**Figure 5F** and **S9**). To elucidate the detailed permeation mechanism of Ca^2+^ through the ACA2 WT and its variants, we conducted well-tempered metadynamics simulations in two independent replicas per system. These simulations yielded free energy barriers and profiles for ion translocation (**Figure S9A-B**). For the WT ACA2 and ACA2ΔCa^2+^ (VEND-AAAA) systems, both replicas converged on nearly identical free-energy landscapes and ion pathways, with a maximum variation of 3 kcal mol^−1^, demonstrating reproducibility. In contrast, while the overall shape of the free-energy surface (FES) and the permeation pathway remained consistent between replicates for the WT 2Ca^2+^ systems (**Figure S9**), the barrier heights differ by up to 25 kcal mol^−1^. The Ca^2+^ trajectory from VEND binding site to ER lumen in ACA2 simulation illustrates how the transmembrane helices adopt an E2P-like conformation to facilitate ion passage toward the ER lumen (**Figure 5G**). Compared to the initial E1P-like starting structure (**Figure. 5H**), the pore exhibits a markedly wider opening during permeation. Among the transmembrane segments, helices M1–M5 undergo the largest displacements, whereas M6, which forms the core of the pore, remains relatively rigid. This helix rearrangement, involving opening of the luminal gate helices M1-M6 and subsequent Ca^2+^ release, mirrors the gating mechanism described for other P-type Ca^2+^-ATPases (Kobayashi et al., 2021). The total combined inter-helix distances between helices M1–M6 and M2–M4 were further analysed, revealing a distinct pattern between WT and the mutants (**Fig. S9C**). Whereas the ACA2 opens its luminal gate gradually, ACA2ΔCa^2+^ (VEND-AAAA mutant) requires much less energy for Ca^2+^ to traverse the pore, following a more “permissive” gating profile (**Figures 5F and S9**). In the ACA2ΔCa^2+^ (VEND-AAAA mutant), disruption of the Ca^2+^-binding site lowers the energy barrier by ∼ 30 kcal mol^−1^ relative to ACA2, and the luminal gate does not need to open fully to allow ion passage, greatly facilitating permeation. Loss of this binding site abolishes the requirement to open the luminal gate (**Figure S9**), suggesting that VEND-AAAA converts the transporter into a more permissive, less selective, and possibly “leaky” channel. To test this hypothesis, we replaced Ca^2+^ with Mg^2+^ to assess ACA2 permeability with a modified Ca^2+^-binding site (**Figure 5F** and **Figure S9**). The energy required to transport Mg^2+^ is similar to that for Ca^2+^, further supporting the idea that this modification makes the ACA2 less selective, resulting in a leakier conduction pathway. The ACA2 WT 2Ca^2+^ system, although not a mutant, required higher energy for Ca^2+^ permeation (**Figure 5F**). This increased barrier arises because the transmembrane helices in this system have greater difficulty rearranging into an open conformation (**Figures. S9**). These observations suggest that the second Ca^2+^ ion placed in the DKTGT phosphorylation motif promotes an occluded state of the transmembrane domain by mimicking the E1P state through the bridging of the P and N domains, triggering the A-domain to “push” on the M1–M6 helices, thereby hindering the opening of the luminal gate and occluding the luminal exit. Overall, all these results together help us to propose a working model of ACA2 action in growing RH. Our findings suggest that Ca^2+^ is transported to internal reservoirs, to maintain Ca^2+^ gradient throughout RH development, and emphasize the ER function in modulating Ca^2+^_cyt_ dynamics during RH growth (**Figure 6**). Since the level of identity between ACA2 and ACA7 protein sequences is extremely high (92.9%) with all key residues in ER-location and Ca^2+^ binding conserved (**Figure S10**), we suggest that results obtained here for ACA2 could be extrapolated to ACA7 protein.

**Figure 5.**
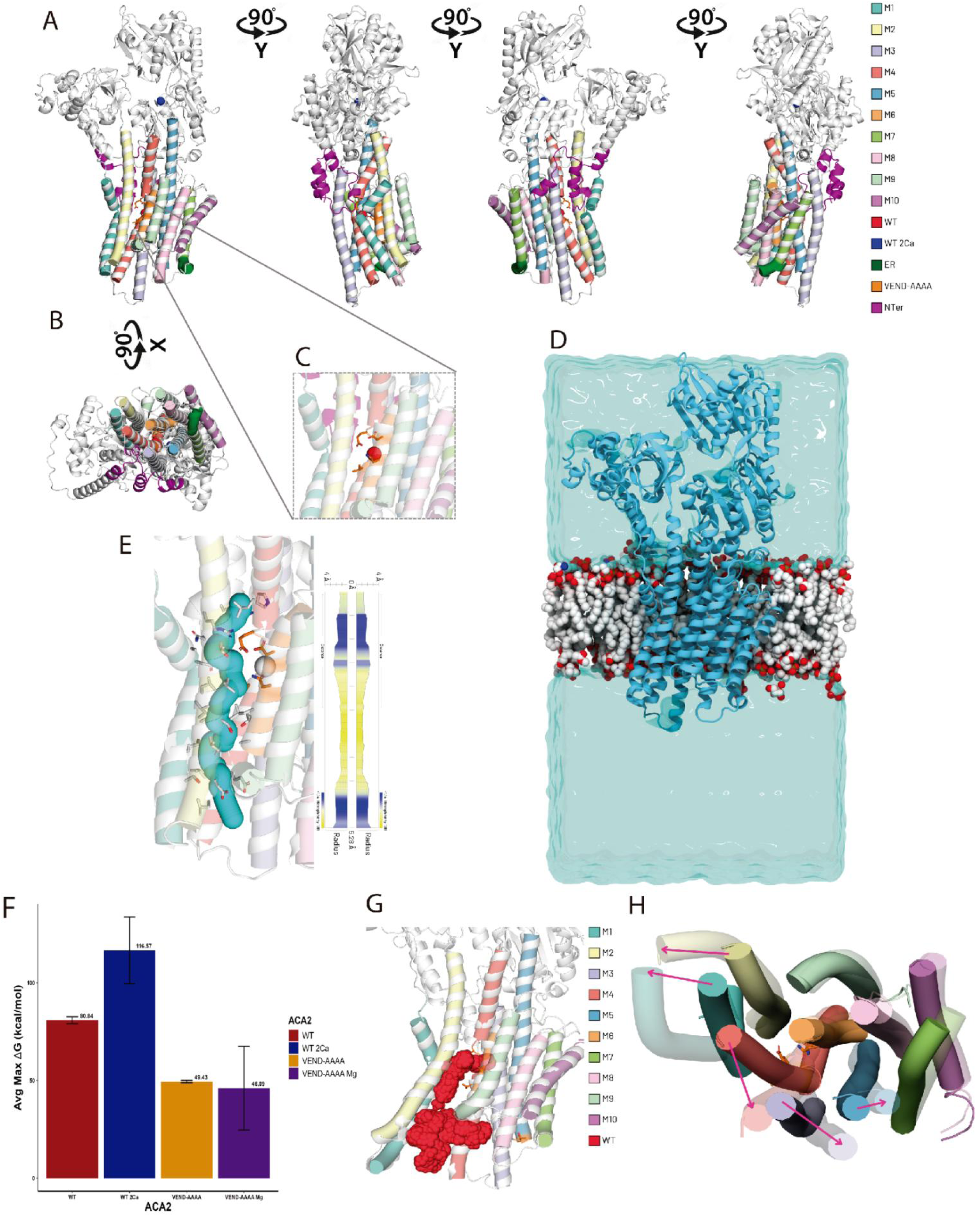
Three-dimensional structure of ACA2 and its variants. **Ca^2+^ and Mg**^2+^ **permeation mechanism through ACA2 variants.** (**A**) Panoramic view of ACA2. Transmembrane helices M1–M10 are shown as colored cylinders, with each variant’s altered segment highlighted. (**B**) Bottom view of the ACA2 transmembrane domain. (**C**) Atomic detail of the Ca^2+^-binding site for ACA2ΔCa^2+^: VEND residues, which are mutated to AAAA in the VEND-AAAA variant, are shown. (**D**) ACA2 embedded into endoplasmic reticulum membrane and solvated in water. (**E**) Luminal pore region with adjacent residues. Hydrophobicity of pore-lining surfaces is indicated. (**F**) Relative energy barriers for Ca^2+^ and Mg^2+^ permeation through the ACA2 pore, averaged across replicas. Standard deviations are indicated by error bars. (**G**) Ca^2+^ trajectory through ACA2 WT from the VEND binding site to the ER lumen. (**H**) Luminal gate opening during ACA2 WT meta-dynamics simulation. Opaque helices represent the initial pore conformation, while transparent helices show their positions upon gate opening. Pink arrows highlight the helices that underwent the most significant movement.

**Figure 6.**
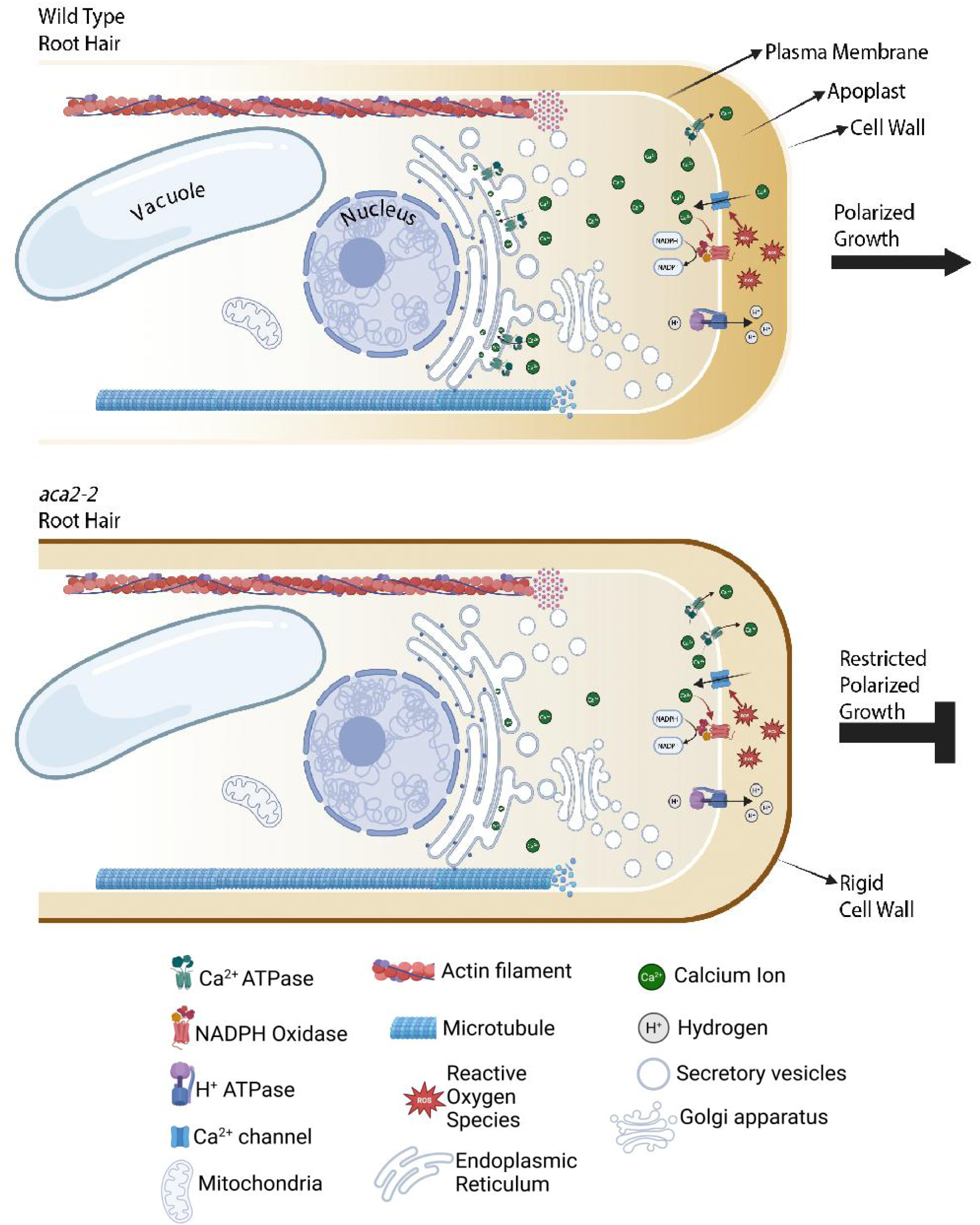
Proposed model of ACA2 action in growing RH. Our findings align with a concept in which Ca^2+^ is transported to internal reservoirs, hence regulating the establishment and maintenance of the Ca^2+^ gradient throughout RH development, and underscore the critical function of the ER organelle in the modulation of Ca^2+^_cyt_ dynamics during RH growth. Changed dynamics in Ca^2+^_cyt_ levels may trigger putative higher Ca^2+^_apo_ leading to the excessive rigidification of cell walls via enhanced “egg box” formation. The alterations in cell wall structures may influence the process of cell elongation. Our results might integrate into a model in which the influx of ions from the apoplast to the cytosol is regulated by a network of Ca^2+^-permeable channels (e.g., GLRs, MSLs, OSCAs, and CNGCs) situated at the apex of the RH. Created at https://BioRender.com.

### Conclusions

The results of our research indicate that the lack of RH ACA proteins, particularly those situated in the ER (ACA2 and ACA7), causes reduced RH growth. This agrees with the evidence that members of the ACA1, ACA2 and ACA7 subgroup P2B-1 are functionally interchangeable and predominantly located in the ER (Rahmati Ishka et al., 2021). These authors also showed that the failure of all three pumps (*aca1/aca2/aca7*) led to plants exhibiting various phenotypes, such as deficits in pollen transmission and SA-dependent lesions in rosette leaves while Ca^2+^_cyt_ imaging in leaves revealed that *aca1/aca2/aca7* mutants exhibit Ca^2+^ transients characterized by heightened magnitudes and prolonged durations upon stimulation. These findings together with our work underscore the significance of the ER-localized ACAs subgroup in regulating cellular Ca^2+^ dynamics and imply that variations in their ACA activities may influence various aspects of plant growth and environmental responses. However, the triple mutant *aca1/aca2/aca7* showed an increase in Ca^2+^ cyt levels either when stimulated with the pathogen elicitor flg22 or under blue-light stimulus (Rahmati Ishka et al., 2021). Our study differs from what we have observed recently for pollen tube growth (García Bossi et al., 2026) where the vacuolar ACAs (ACA4 and ACA11) have more significant effect than the ER-ACAs (ACA2 and ACA7) in the cell elongation process linked to the regulation of Ca^2+^_cyt_ dynamics. This may indicate that although both plant cell types grow in a similar polar manner, they are not exactly comparable. The decreased [Ca^2+^]_cyt_ levels detected in *aca2-2* are associated with a deficiency in RH elongation. Our results align with a model where Ca^2+^ is conveyed to internal reservoirs, regulating the formation and maintenance of the Ca^2+^ gradient throughout RH growth. It highlights a significant role of ER organelle in the regulation of [Ca^2+^]_cyt_ and dynamics during RH growth. Whereas in stomatal guard cells a proper ER Ca^2+^ homeostasis is required to maintain the spontaneous Ca^2+^_cyt_ oscillations and regulate stomatal aperture (Grenzi et al., 2025) the impact of a deregulated ER Ca^2+^ transport is not known. Our hypothesis for RH is that ACA2 participates in the feedback loop regulating [Ca^2+^]_cyt_ dynamics and determining the homeostatic setpoint. [Ca^2+^]_cyt_ oscillations in *aca2-2* tend to be slower and more regular, with a lower baseline. Having longer periods of Ca^2+^ oscillations, as well as a lower cytosolic Ca^2+^ concentration in the root hair tip in *aca2-2* may affect the dynamic discharge of ions from the cytosol to the ER impacting the correct induction of RH elongation. Our findings are incorporated into a proposed model (**Figure 6**) whereby the inflow of ions from the apoplast to the cytosol is modulated by a network of Ca^2+^-permeable channels (e.g. GLRs, MSLs, OSCAs, and CNGCs) located at the apex of the RH. To return Ca^2+^_cyt_ to its baseline levels, Ca^2+^ would be translocated to the apoplast via plasma membrane ACAs (like ACA8/9/10/12/13), and/or to organelles by ACA pumps (examined herein) or CAX exchangers. Additional research is necessary to elucidate the interplay between ACAs, CAXs, and influx channels in sustaining cytosolic Ca^2+^ homeostasis during both resting and activated states. The precise roles of other Ca^2+^ transporters, such as ECAs, are still unclear. Further research is required to ascertain if the ACA pumps only facilitate the influx of Ca^2+^ into organelles or whether they also enable the efflux from organelles to the cytosol. Our findings illustrate the importance of ACA2 in sustaining the Ca^2+^_cyt_ gradient during RH growth. Finally, one major technical limitation to our study is the difficulties in measuring the dynamics of Ca^2+^_ER_ and in Ca^2+^_apo_ in growing RHs.

## Materials and methods

### *In silico* gene expression analysis

A preliminary search of the different ACAs (ACA1/AT1G27770, ACA2/AT4G37640, ACA4/AT2G41560, ACA7/AT2G22950, ACA8/AT5G57110, ACA9/AT3G21180, ACA10/AT4G29900, ACA11/AT3G57330, ACA12/AT3G63380, ACA13/AT3G22910) was made at the Root Cell Atlas (https://rootcellatlas.org/) to identify which ones had a higher expression in the epidermis and in particular in trichoblasts. This atlas uses the Seurat toolkit to integrate the preprocessed data from seven different studies (Denyer et al., 2019; Jean-Baptiste et al., 2019; Ryu et al., 2019; Shahan et al., 2022; Shulse et al., 2019; Wendrich et al., 2020; T.-Q. Zhang et al., 2019). A second search to integrate the relative levels of expression was made at a different atlas (Shahan et al., 2022). For this analysis, only ACAs that showed expression in the root were included.

### Growth conditions

All plant materials used in this study were in the Columbia-0 ecotype background of *Arabidopsis thaliana*. Seeds were sterilized and placed on half-strength (½MS) Murashige and Skoog (MS) medium (Duchefa) pH 5.8 supplemented with 0.8% agar. For root measurements, RNA extraction and confocal microscopy 7-day-old seedlings were grown on square plates placed vertically at 22°C with continuous light, after stratification in dark at 4°C for 1 d on the plates. Seedlings on plates were transferred to soil and kept in the greenhouse in long-day conditions to obtain mature plants for transformation, genetic crossing, and amplification of seeds.

### Plant material

For identification of homozygous T-DNA knockout lines, genomic DNA was extracted from rosette leaves. Confirmation by PCR of a unique band corresponding to T-DNA insertion in the target genes ACA2 (AT4G37640: SALK_082624 (*aca2-1*); SAIL_873_F03 (*aca2-2*)), ACA4 (AT2G41560: SALK_029620 (*aca4-1*); SALK_002135 (*aca4-2*)), ACA7 (AT2G22950: SALK_014124 (*aca7-1*); SALK_143504 (*aca7-2*)) and ACA11 (AT3G57330: SAIL_269_C07 (*aca11-1*); SALK121482 (*aca11-2*)) were performed using an insertion-specific LBb1.3 for SALK lines or Lb1 for SAIL lines. Mutant and transgenic lines are listed in **Table S1** and primers used are listed in Supporting Information **Table S2**.

### RH length measurement

To determine the RH length, images of root tips were taken using an Olympus stereo microscope SZX7 at 32x magnification. RH length was measured using ImageJ at the maturation zone, measuring 10 RH per plant.

### Statistical analysis

The statistical analysis was performed using the R statistical software (R Development Core Team., Retrieved March 17, 2025) through the integrated development environment RStudio (Posit Team, 2025). The package “nlme” (Bates & R Core Team, 2025) was employed for Generalized Least Squares (GLS). Tukey’s multiple comparisons (post hoc tests) were conducted using the “emmeans” (Lenth, 2024) and “multcomp” (Hothorn et al., 2008) packages. Additionally, the “car” (Fox & Weisberg, 2011) package was used to fit ANOVA tables.

### ACA2 variants

Synthetic DNA was designed containing full length ACA2 and ACA2ΔCa^2+^ cDNA containing GatewayTM (Life Technologies, Carlsbad, CA, USA) attB1 and attB2 sites. Recombinase-mediated integration of the PCR fragment was made into pDONR201 (Invitrogen). pDONR201/ACA2 and pDONR201/ ACA2ΔCa^2+^ constructions were recombined into the vector pGWB405 (Kanamycin R, Invitrogen, Carlsbad, CA, USA) to overexpress ACA2 and ACA2ΔCa^2+^ under 35S mosaic virus promoter and add a GFP tag to the C-terminal of the protein (35S::ACA2-GFP and 35s:: ACA2ΔCa^2+^-GFP). Wild-type and T-DNA *aca2-2* mutant plants were transformed by using Agrobacterium (strain GV3101+pSoup). Plants were selected with kanamycin (50 μg/ml) and several independent transgenic plants were isolated for each construct.

### Transient expression assays in *Nicotiana benthamiana*

To test the sub-cellular localization of ACA2 and variants, Four-week-old *Nicotiana benthamiana* leaves were infiltrated with Agrobacterium strains (GV3101) carrying either 35S::ACA2-GFP, 35s:: ACA2ΔER-GFP, or 35s:: ACA2ΔCa^2+^-GFP and ER-KDEL-Cherry constructs. After 2 d, images of the lower leaf epidermal cells were taken using a confocal microscope (Zeiss LSM 710).

### COS-488 staining

After surface sterilization in 70% ethanol, seeds were vernalized at 4°C for 2 days in darkness, afterwards grown vertically on plates with 1⁄2 strength Murashige and Skoog (MS) without sugar medium containing 1% sucrose in a long-day regime (16 h light and 8 h darkness) at 21°C. For image acquisition, an upright Leica TCS SP8 FALCON FLIM confocal laser scanning microscope, equipped with a Leica HC PL APO Corr 63 x 1.20 water immersion CS2 objective, was used. AlexaFluor-COS 488 was excited at 488 nm (fluorescence emission: 500 - 550 nm). The signal intensity of intracellular space or plasma membrane was assessed using Fiji. The COS probe was generated as described in Mravec *et al.,* 2014, using chitosan oligosaccharides conjugated with AlexaFluor 488 hydroxylamine (ThermoFisher, Waltham, Massachusetts, USA). In brief, 1 mg/ml oligosaccharide solution was mixed into 0.1 M sodium acetate buffer, pH 4.9. The AlexaFluor 488 (10 mg/ml in DMSO) was added to the oligosaccharide (1⁄2 of the moles of the oligosaccharide). After incubation at 37°C for 48 h with shaking at 1,400 rpm the probe was ready. 6-day-old seedlings were stained in liquid 1⁄2 MS medium without sugar supplemented with COS488 at a 1:500 dilution, for 30 min, and washed with medium. Imaging root tips was performed directly afterwards.

### Confocal microscopy

Confocal laser scanning microscopy was performed using Zeiss LSM 710 and LSM5 Pascal. ACA2 variants: Fluorescence was analysed by using a laser line of 488 nm for GFP excitation and emitted fluorescence was recorded between 490 and 525 nm. Images acquired with W Plan-APOCHROMAT 20x objective (N/A 1.0). Transient expression in Nicotiana: Fluorescence was analysed by using a laser line of 488 nm for GFP and 543 nm for mCherry excitation and emitted fluorescence was recorded between 490 and 525 nm for GFP and 590 and 650 nm for mCherry. Images acquired with W Plan-APOCHROMAT 20x objective (N/A 1.0). For colocalization quantification between fluorophores, we used ZEN 2.3 SP1 software (Zeiss), where the “Colocalization” tool calculates the overlap between channels using Manders coefficient (Manders et al., 1993). The Manders coefficient is reported as M1, defined as the fraction of the GFP (ACA2 variant) signal that overlaps with the mCherry (ER-KDEL) channel.

### Light Sheet Fluorescence Microscopy (LSFM)

LSFM was performed in transgenic *Arabidopsis thaliana* (Col-0 and insertional mutants previously mentioned) expressing the cytosolic-targeted FRET based sensor Cameleon NES-YC3.6 (Krebs et al., 2012). We used the set up for FRET measurement described in (Candeo et al., 2017). Here we provide a brief description of sample preparation and imaging methods, for an in-depth description of these protocols we recommend referring to (Romano Armada et al., 2019)). Sample preparation and seedling transfer were performed under sterile conditions, in a laminar flow. Sample tube preparation: Fluorinated Ethylene Propylene (FEP) with internal diameter of 2 mm were cut into 3 cm pieces and assembled into the crown of 10 µl pipette tips. The tubes were sonicated with NaOH 0.5 M for 5 min and with ethanol 70% for 5 min as well. Then they were rinsed 3 times with distilled water, placed in a pipette tip box, left to dry and autoclaved at 121°C for 15 min. Sample preparation: Seeds were sterilized with 70% ethanol for 15 min and rinsed twice with 100% ethanol. Seeds were sown in Petri dishes with ½MS medium pH 5.8 supplemented with 0.8% plant agar. The plates were placed at 4°C for 2 d for seed stratification. Later they were transferred to a growth chamber with 16/8h light/dark cycle at 22°C for 2 d for seed germination. Seedling transfer: FEP tubes were filled with melted ½ MS media supplemented with 0.5% Gelrite^TM^ (Duchefa) and left to dry in horizontal position. The upper part of the tube was sealed with melted ½ MS media supplemented with 0.8% plant agar and tubes were placed once again into the tip box. Two-day-old seedlings were transferred to the tubes, and the tip box was filled with ½ MS liquid media to prevent the media in the tubes from drying. Boxes were placed in a growth chamber with 16/8h light/dark cycle at 22°C for 5 d. Image acquisition: For the acquisition, the parameters used were set differently to the 10x and 20x magnification, respectively, as follows: 270 and 400 time points (each time point acquisition) every 2.5 s and 3 s, with an exposure time of 0.1 s for both. The number of acquired planes were N=12 and N=15 (sections scanned with the light sheet at each time point) every Δz = 4 µm (linear step between planes). For the 10x dataset, data processing was performed using Napari software. The tip of each RH was selected using the napari-roi-registration plug in (available on the Napari hub - https://www.napari-hub.org/plugins/napari-roi-registration), allowing us to track RH growth as well as the changes in fluorescence intensity. For the 20x dataset, image analysis was performed using the Fiji plugin ‘Template Matching’ (https://sites.google.com/site/qingzongtseng/template-matching-ij-plugin) after applying a Gaussian blur filter with 0.7 sigma, selecting the tip of each RH as a landmark and the subpixel cross-correlation method. Both methods yielded results that allowed us to track changes in RH growth and fluorescence intensity.

### Phenotyping oscillations

Initially, an overall analysis of the [Ca^2+^]_cyt_ oscillations was performed with Fourier transforms for each time series of both genotypes (Col-0 and *aca2-2*), using the built-in MATLAB function and yielding two average Power Spectra Distribution. As described in Candeo et al. (2017), power peaks were indicated as ‘High’ (HF) and ‘Low Frequency’ (LF) components. To detect possible phenotypes in oscillations, a pipeline in R statistical software (R Development Core Team., Retrieved March 17, 2025) was tailored to preprocess individual time series, followed by wavelet analysis and ending in population-level comparisons. The synchronization between pairs of time series was also evaluated with a cross-wavelet analysis, here between RH growth rate and the fluorescence ratio (cpVenus/CFP) of the YC3.6 Ca^2+^Regression Fitting (loess) to extract trend and spurious lower frequency components, employing high span values (>0.15 and of degree 2). The resulting trace included trend and the LF component, which was separately analyzed. Growth time series were extracted from smoothing the position series and then differentiating, which resulted in the instantaneous growth rate. Oscillation analysis was based on the Continuous Wavelet Transform from the ‘WaveletComp’ package (Roesch & Regression Fitting (loess) to extract trend and spurious lower frequency components, employing high span values (>0.15 and of degree 2). The resulting trace included trend and the LF component, which was separately analyzed. Growth time series were extracted from smoothing the position series and then differentiating, which resulted in the instantaneous growth rate. Oscillation analysis was based on the Continuous Wavelet Transform from the ‘WaveletComp’ package (Roesch & Schmidbauer, 2025), using the Morlet wavelet to assign a power value to different periods in each time point. The significance of these time explicit components in the wavelet spectrum was assessed using a null model of an Autoregressive process of order 1 (AR(1)), a red noise test based in Torrence & Compo (1998). We considered only the periods whose p-value < 0.05 and were outside the Cone of Influence, shaded area in the wavelet spectra corresponding to a region with considerable edge effects of the analysis. The main periodic component was extracted from the period with the highest power in each time point, being retrieved along with the respective power and phase values for individual and pairs of time series. This enables us to investigate not only the overall oscillatory behavior of the series, but also its stability across time. Measures of centrality (e.g. mean and median) and dispersion (e.g. standard deviation and coefficient of variation) were obtained for all the variables, allowing comparisons between Col-0 and *aca2-2* mutant. Regression Fitting (loess) to extract trend and spurious lower frequency components, employing high span values (>0.15 and of degree 2). The resulting trace included trend and the LF component, which was separately analyzed. Growth time series were extracted from smoothing the position series and then differentiating, which resulted in the instantaneous growth rate. Oscillation analysis was based on the Continuous Wavelet Transform from the ‘WaveletComp’ package (Roesch & Schmidbauer, 2025), using the Morlet wavelet to assign a power value to different periods in each time point. The significance of these time explicit components in the wavelet spectrum was assessed using a null model of an Autoregressive process of order 1 (AR(1)), a red noise test based in Torrence & Compo (1998). We considered only the periods whose p-value < 0.05 and were outside the Cone of Influence, shaded area in the wavelet spectra corresponding to a region with considerable edge effects of the analysis. The main periodic component was extracted from the period with the highest power in each time point, being retrieved along with the respective power and phase values for individual and pairs of time series. This enables us to investigate not only the overall oscillatory behavior of the series, but also its stability across time. Measures of centrality (e.g. mean and median) and dispersion (e.g. standard deviation and coefficient of variation) were obtained for all the variables, allowing comparisons between Col-0 and *aca2-2* mutant, using the Morlet wavelet to assign a power value to different periods in each time point. The significance of these time explicit components in the wavelet spectrum was assessed using a null model of an Autoregressive process of order 1 (AR(1)), a red noise test based in Torrence & Compo (1998). We considered only the periods whose p-value < 0.05 and were outside the Cone of Influence, shaded area in the wavelet spectra corresponding to a region with considerable edge effects of the analysis. The main periodic component was extracted from the period with the highest power in each time point, being retrieved along with the respective power and phase values for individual and pairs of time series. This enables us to investigate not only the overall oscillatory behavior of the series, but also its stability across time. Measures of centrality (e.g. mean and median) and dispersion (e.g. standard deviation and coefficient of variation) were obtained for all the variables, allowing comparisons between Col-0 and Regression Fitting (loess) to extract trend and spurious lower frequency components, employing high span values (>0.15 and of degree 2). The resulting trace included trend and the LF component, which was separately analyzed. Growth time series were extracted from smoothing the position series and then differentiating, which resulted in the instantaneous growth rate. Oscillation analysis was based on the Continuous Wavelet Transform from the ‘WaveletComp’ package (Roesch & Schmidbauer, 2025), using the Morlet wavelet to assign a power value to different periods in each time point. The significance of these time explicit components in the wavelet spectrum was assessed using a null model of an Autoregressive process of order 1 (AR(1)), a red noise test based in Torrence & Compo (1998). We considered only the periods whose p-value < 0.05 and were outside the Cone of Influence, shaded area in the wavelet spectra corresponding to a region with considerable edge effects of the analysis. The main periodic component was extracted from the period with the highest power in each time point, being retrieved along with the respective power and phase values for individual and pairs of time series. This enables us to investigate not only the overall oscillatory behavior of the series, but also its stability across time. Measures of centrality (e.g. mean and median) and dispersion (e.g. standard deviation and coefficient of variation) were obtained for all the variables, allowing comparisons between Col-0 and *aca2-2* mutant.

### Molecular Modeling of ACA2

The structures of the ACA2 WT and its mutant variants were modeled using AlphaFold3 based on their amino acid sequences (Abramson et al., 2024). In total, four ACA2 variants were generated. The ACA2 WT with the unmodified protein, with one Ca^2+^ ion placed inside the pore and coordinated by residues V407, E412, N849, and D853 and ACA2 WT 2Ca with an additional Ca^2+^ ion placed in the phosphorylation motif DKTGT. ACA2 VEND-AAAA in which the Ca^2+^–binding site VEND was substituted with AAAA and ACA2 VEND-AAAA Mg^2+^ in which the Ca^2+^ was replaced with a Mg^2+^ ion. Each structure was processed with CHARMM-GUI (Jo et al., 2008), and system topologies were built using the CHARMM36m force field for GROMACS (Huang et al., 2017; J. Lee et al., 2016). Ca^2+^ ions were described with the multi-site Ca2+ model optimized for Ca2+–protein interactions, which accurately represents the permeation mechanism through channels (A. Zhang et al., 2020). Proteins were embedded in a mixed lipid bilayer composed of palmitoyl-linoleoyl phosphatidylcholine (PLPC, 55%), stearoyl-linoleoyl phosphatidylethanolamine (SLPE, 30%), palmitoyl-linoleoyl phosphatidic acid (PLPA, 5%), palmitoyl-linoleoyl phosphatidylinositol (PLPI, 5%), and palmitoyl-linoleoyl phosphatidylserine (PLPS, 5%) to mimic the *A. thaliana* ER membrane (Bahammou et al., 2024; Reszczyńska & Hanaka, 2020). MD simulations were performed with GROMACS 2024 (Abraham et al., 2015), following an energy minimization and equilibration protocol adapted for protein–membrane systems (Cabral et al., 2024; Villavicencio et al., 2022). Briefly, each system underwent: (1) Energy minimization using the steepest-descent algorithm until the step size becomes too small, indicating that the system energy can no longer be reduced; (2) One 1 ns NVT step with position restraints of 5,000 kJ mol^−1^ on protein heavy atoms and 1,000 kJ mol^−1^ on lipid heavy atoms; (3) Six successive 1 ns NPT steps with progressively relaxed restraints: the first NPT step retained the same force constants as in NVT; each subsequent step reduced protein restraints by 1,000 kJ mol^−1^ and lipid restraints by 200 kJ mol^−1^. The final NPT step applied a 1,000 kJ mol^−1^ restraint only to protein backbone heavy atoms. Production runs (metadynamics) with all position restraints removed. Throughout the simulations, a 2 fs time step was used with the LINCS algorithm to constrain hydrogen bonds (Hess, 2008). The V-rescale thermostat (relaxation time τ = 0.1 ps) and the C-rescale barostat (τ = 2 ps) maintained temperature and pressure (Bernetti & Bussi, 2020; Bussi et al., 2007). Nonbonded interactions employed a 1.2 nm cutoff and the Particle Mesh Ewald method for long-range electrostatics (Darden et al., 1993; Gonçalves et al., 2019). For production simulations, the Parrinello–Rahman barostat was used (τ = 2 ps) with no position restraints (Parrinello & Rahman, 1981).

### Metadynamics of ion permeation in ACA2

To investigate ion permeation through the ACA2 pore in all five systems, it was performed with well-tempered metadynamics using PLUMED 2.9 (Barducci et al., 2008; Tribello et al., 2014). For each system, the starting structure was the final frame from equilibration. We biased two collective variables (CVs) that describe the geometry of the luminal gate forming helices M1, M2, M4, and M6. CV1: the distance between the center of mass (COM) of the ion and the COM of residues T191, K196, S388, and A842, which lie at the luminal termini of helices M1, M2, M4, and M6, respectively. CV2: the sum (x + y) of the COM distance between helices M1 and M6 (x) and the COM distance between helices M2 and M4 (y). Both CVs were biased with Gaussian potentials of height 1.0 kJ mol−1, a bias factor of 20, and a deposition interval of 200 MD steps. Gaussian widths were 0.3 nm for CV1 and 0.5 nm for CV2. Two independent replicas were run per system. Simulations were terminated once CV1 reached 0.1 nm, signifying that the ion had traversed the channel. This protocol limits sampling to the pore region and prevents distortion of the free energy surface (FES) by sampling ion positions in the luminal solvent or outside the pore.

## Acknowledgements

We would like to thank Melanie Krebs for kindly providing the NES-YC3.6 construct. We are grateful to Jeff Harper for sharing seed lines and Dr. Andrés Rossi, Dr Alejandra Ross, and Dr Esteban Miglietta, from FIL microscopy facility, for technical support with confocal microscopy. Dr Santiago Mora Garcia for sharing *Nicotiana benthamiana* plants and ER maker, Romina Acha for helping us with purchasing materials and Silvina Paola Denita Juarez for initial Ca^2+^ imaging on YC3.6 in confocal microscopy at FIL. We thank ABRC (Ohio State University) for providing T-DNA seed lines. J.M.E. and J.P.M. are investigators of the National Research Council (CONICET) from Argentina. This work is supported from ANPCyT (PICT2021-0514), by ANID – Programa Iniciativa Científica Milenio ICN17_022, and Fondo Nacional de Desarrollo Científico y Tecnológico [1250304] to J.M.E. A.C. and S.B. acknowledge the Ministero dell’Istruzione, dell’Università e della Ricerca — Fondo per Progetti di Ricerca di Rilevante Interesse Nazionale 2022 (PRIN 2022NMSFHN); A.C. the Agritech National Research Center, funded by the European Union NextGenerationEU (Piano Nazionale di Ripresa e Resilienza (PNRR) — Missione 4, Componente 2, Investimento 1.4 – D.D. 1032 17/06/2022, CN00000022), and S.B. The “Fondazione Fratelli Confalonieri”; M.T.P and A.F.S.M. acknowledge the São Paulo Research Foundation (FAPESP 2019/26129-6; 2025/22641-5).

## Author Contribution

M.C.S. performed most of the experiments, analysed the data and helped with the writing process of the manuscript. V.C and H.V. performed the molecular dynamics characterization of ACA2 variants. J.G.B. helped with the mutant line generation and R-GECO transgenics. S.B., A.C., G.T., helped with LSFM imaging. C.B. helped at the beginning of this project. P.R.M. and E.B. performed COS-488 staining. V.B.G. helped analyse the data. J.M.P. and D.R.R.G. gave feedback on the manuscript. C.M. generated the ACA2 structure and variants. J.P.M. provided mutant and transgenic lines, analysed the data and helped in the writing process. A.B. and A.C helped with the LSFM set up, data acquisitions and data analysis. A.F.S.M., M.T.P., D.S.C.D. performed Ca^2+^ analysis. J.M.E. designed research, analysed the data, supervised the project, and wrote the paper. All authors commented on the results and the manuscript. This manuscript has not been published and is not under consideration for publication elsewhere. All the authors have read the manuscript and have approved this submission.

## Competing financial interest

The authors declare no competing financial interests. Correspondence and requests for materials should be addressed to J.M.E.

## Notes

### Competing Interest Statement

The authors have declared no competing interest.

